# A large effective population size for within-host influenza virus infection

**DOI:** 10.1101/2020.04.17.046250

**Authors:** Casper K Lumby, Lei Zhao, Judy Breuer, Christopher J R Illingworth

## Abstract

Strains of the influenza virus form coherent global populations, yet exist at the level of single infections in individual hosts. The relationship between these scales is a critical topic for understanding viral evolution. Here we investigate the within-host relationship between selection and the stochastic effects of genetic drift, estimating an effective population size of infection N_e_ for influenza infection. Examining whole-genome sequence data describing a chronic case of influenza B in a severely immunocompromised child we infer an N_e_ of 2.5 × 10^7^ (95% confidence range 1.0 × 10^7^ to 9.0 × 10^7^) suggesting the importance of genetic drift to be minimal. Our result, supported by the analysis of data from influenza A infection, suggests that positive selection during within-host infection is primarily limited by the typically short period of infection. Atypically long infections may have a disproportionate influence upon global patterns of viral evolution.

---

The evolution of the influenza virus may be considered across a broad range of scales. On a global level, populations exhibit coherent behaviour^1–3^, evolving rapidly under collective host immune pressure^4,5^. On another level, these global populations are nothing more than very large numbers of individual host infections, separated by transmission events.

Despite the clear role for selection in global influenza populations, recent studies of within-host infection have suggested that positive selection does not strongly influence evolution at this smaller scale^6–8^. Contrasting explanations have been given for this, with suggestions either that selection at the within-host level is intrinsically inefficient, being dominated by stochastic processes^7^, or that while selection is efficient, a mismatch in timing between the peak viral titre and the host adaptive immune response prevents selection from taking effect^8^.

To resolve this issue, we evaluated the relative importance of selection and genetic drift during a case of influenza infection. The balance between these factors is determined by the effective size of the population, denoted N_e_. If N_e_ is high, selection will outweigh genetic drift, even where differences in viral fitness are small^9^. By contrast, if N_e_ is low, less fit viruses are more likely to outcompete their fitter compatriots.

Estimating N_e_ is a difficult task, with a long history of method development in this area^10–12^. A simple measure of N_e_ may be calculated by matching the genetic change in allele frequencies in a population with the changes occurring in an idealised population evolving under genetic drift^13^. However, such estimates are vulnerable to distortion, for example being reduced by the effect of positive selection in a population. Where the global influenza A/H3N2 population is driven by repeated selective sweeps^14–16^ a neutral estimation method suggests a value for N_e_ not much greater than 100^17^. While methods for jointly estimating N_e_ and selection exist, they are limited to systems with only a few loci of interest^18–22^. Non-trivial population structure can also affect estimates^23^; a growing body of evidence supports the existence of such structure in within-host influenza infection^24–27^. While careful experimental techniques can reduce sequencing error^28^, noise from sequencing and unrepresentative sample collection combine^29^, potentially confounding estimates of N_e_ in viral populations^30^. If N_e_ is high, any signal of drift can be obscured by noise.

We here estimate an effective population size for within-host influenza B infection using data collected from a severely immunocompromised host. While the viral load of the infection was not unusual for a hospitalised childhood infection^31^, an absence of cell-mediated immunity led to the persistence of the infection for several months^32^. Given extensive sequence data collected during infection, the reduced role of positive selection, combined with novel methods to account for noise and population structure, enabled an improved inference of N_e_. The large effective size we infer suggests that selection acts in an efficient manner during within-host influenza infection. The influence of positive selection is limited only by the duration of infection.

## Results and Discussion

Viral samples from the population were collected at 41 time points spanning 8 months during the course of an influenza B infection in a severely immunocompromised host (Fig. 1A). Clinical details of the case, and the use of viral sequence data in evaluating the effectiveness of clinical intervention, have been described elsewhere^32^. After unsuccessful treatment with oseltamivir, zanamivir and nitazoxanide, a bone marrow transplant and favipiravir combination therapy led to the apparent clearance of infection. Apart from a single exception, biweekly samples tested negative for influenza across a period of close to two months. A subsequent resurgence of zanamivir-resistant infection was cleared by favipiravir and zanamivir in combination.

**Figure 1.**
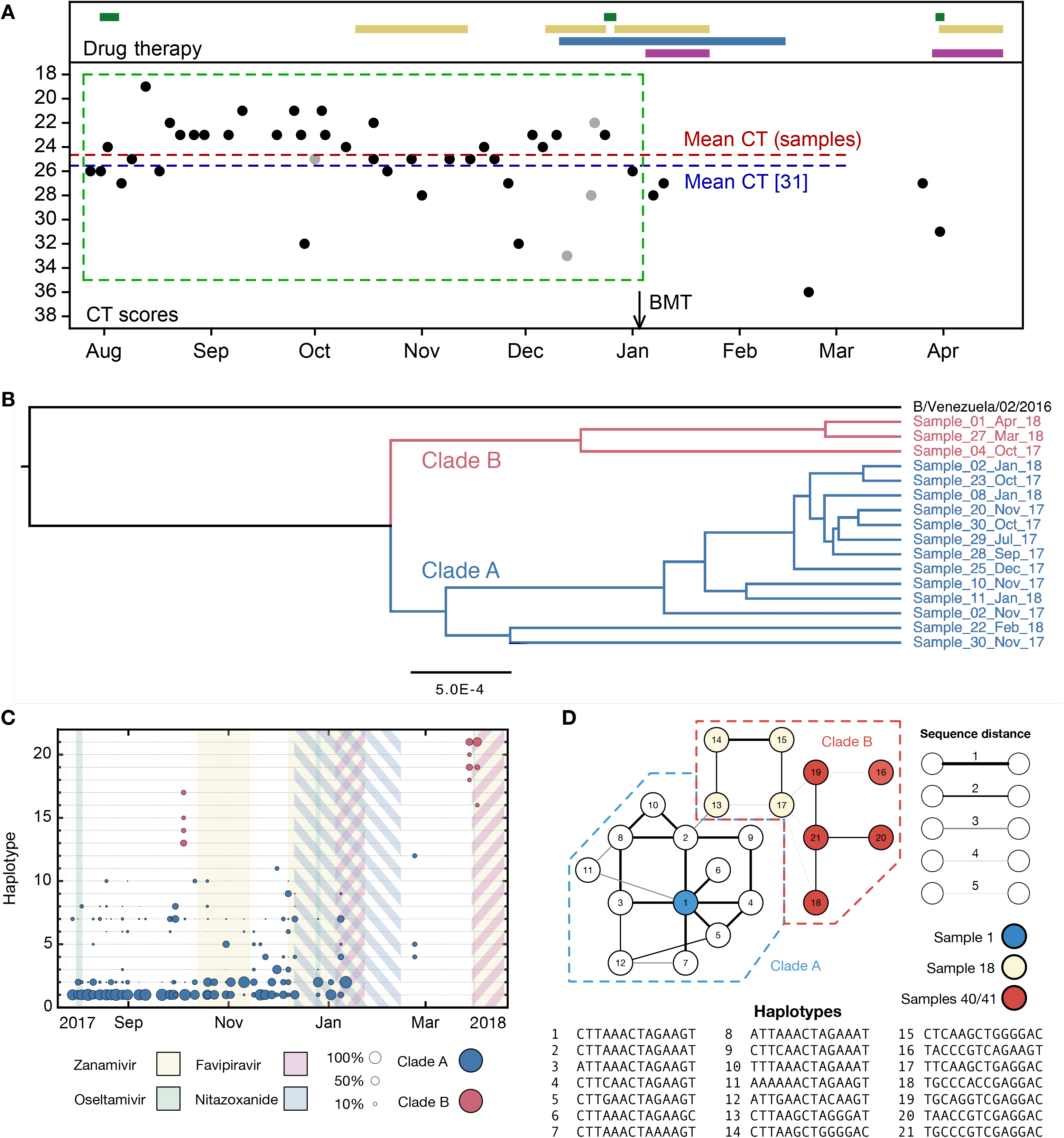
Population structure of the influenza infection. **A.** CT values from viral samples collected over time indicate the viral load of the infection; a higher number corresponds to a lower viral load. Drug information, above, shows the times during which oseltamivir (green), zanamivir (yellow), nitazoxanide (blue) and favipiravir (purple) were prescribed. Black dots show samples from which viral sequence data were collected; gray dots show samples from which viral sequence data were not collected. The green box shows the window of time over which samples were analysed, preceding the use of favipiravir in January. The mean viral load (dashed horizontal line, red) was close to the mean reported for a set of samples from hospitalised children with influenza (dashed horizontal blue line)^31^. A black arrow shows the date of a bone marrow transplant (BMT). **B.** A phylogeny of whole-genome viral consensus sequences identified two distinct clades in the viral population. Clade B featured three samples, distributed across the period of infection, with the remaining samples contained in Clade A. **C.** Sub-consensus structure of the viral population inferred via a haplotype reconstruction algorithm using data from the neuraminidase segment. The same division of sequences into two clades is visible, with samples being comprised of distinct viral genotypes. The area of each circle is proportional to the inferred frequency of the corresponding haplotype in the viral population. Haplotypes reaching a frequency of at least 10% in at least one time point are shown. Multiple drugs were administered to the patient through time, with a favipiravir/zanamivir combination first causing a temporary reduction of the population to undetectable levels, then finally clearing the infection. Haplotypes spanned the loci 96, 170, 177, 402, 403, 483, 571, 653, 968, 973, 1011, 1079, 1170, and 1240 in the NA segment. **D.** Evolutionary relationship between the haplotypes; clade B is distinct from and evolves away from those sequences comprising the initial infection. Numbers refer to the distinct haplotypes identified within the population.

Phylogenetic analysis of whole-genome viral consensus sequences showed the existence of non-trivial population structure, with at least two distinct clades (Fig. 1B, Fig. 1S1); we term these clades A and B. While being phylogenetically separated the two clades persisted across several months of infection. Haplotype reconstruction showed that samples from clade B were comprised of distinct viral haplotypes to those from clade A (Fig. 1C, 1S2). Clade B slowly evolved away from the initial consensus sequence (Fig. 1D), while viruses in clade A stayed close to the initial consensus. The cladal structure suggests the existence of spatially distinct viral populations in the host, samples stochastically representing one population or the other.

To estimate the effective population size, we analysed genome-wide sequence data from samples in clade A collected before first use of favipiravir. A method of linear regression was used to quantify the rate of viral evolution, measuring the genetic distance between samples as a function of increasing time between dates of sample collection. We inferred a rate of 0.051 substitutions per day (97.5% confidence interval 0.034 to 0.068) (Fig. 2A), equivalent to 7.94 substitutions genome-wide across 157 days of evolution. The vertical intercept of this line provides an estimate of the contribution of noise to the measured distance between samples, for example arising from sequencing error or undiagnosed population structure. The identified value of close to 40 substitutions is equivalent to a between-sample allele frequency difference of approximately +/− 0.3% per locus. While considerable noise affects each sample, the dataset as a whole provides a clear signal of evolutionary change.

**Figure 2:**
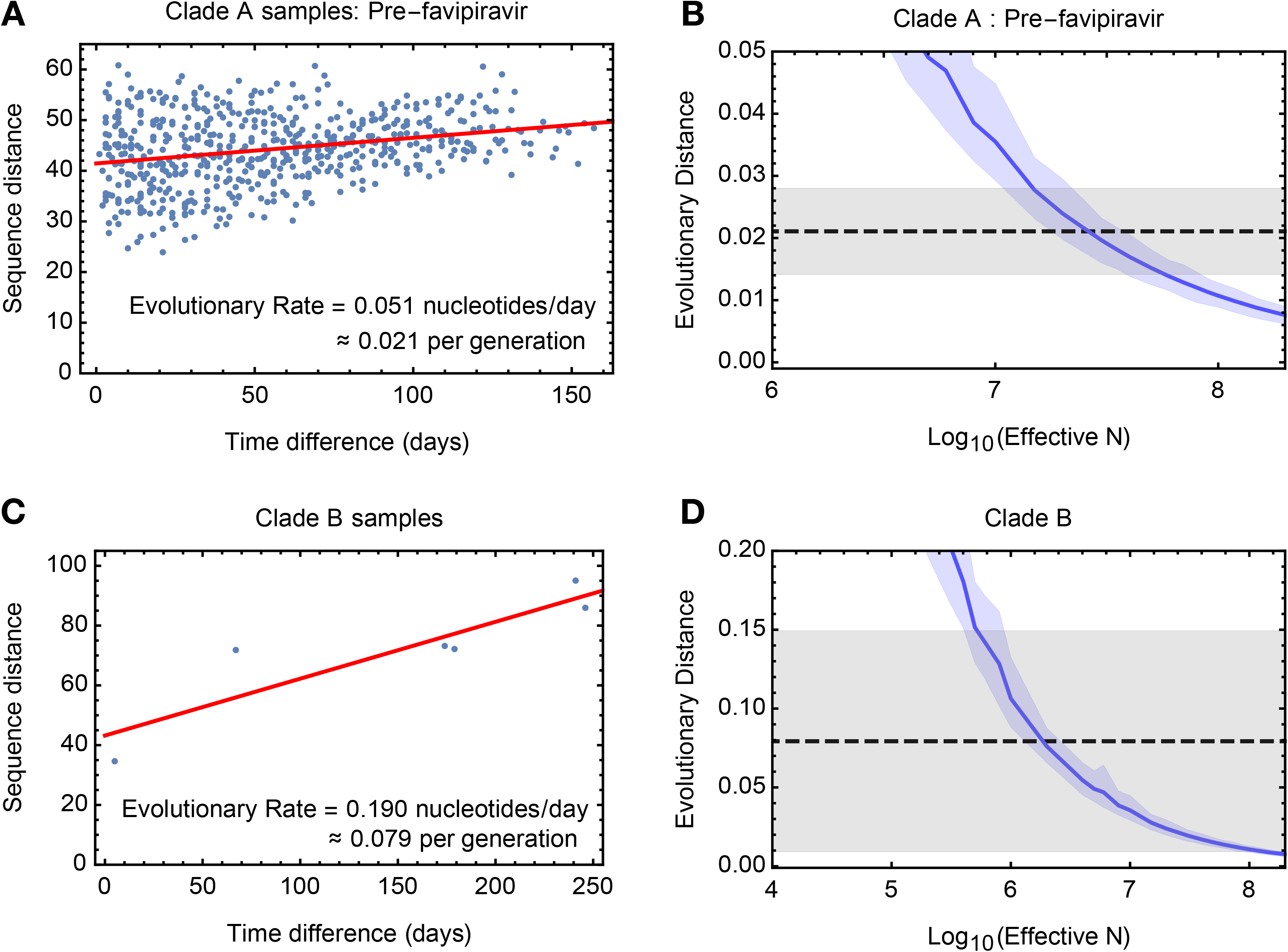
Measuring rates of evolution in the viral population. **A.** Computed rate of evolution for viruses in clade A up to the time of the first use of favipiravir. The distance between two sequences is calculated as the total absolute difference in four-allele frequencies measured across the genome. The calculated rate per generation is based upon a generation time for influenza of 10 hours^53^. **B.** Distribution of evolutionary distances in influenza populations simulated using a Wright-Fisher model compared to the distance per generation calculated in the regression fit. A solid blue line shows the mean, with shading indicating an approximate 97.5% confidence interval around the mean. Statistics were calculated from sets of 400 simulations conducted at each value of N_e_. The dashed black line shows the rate of evolution of the real population; gray shading shows a 97.5% confidence interval for this statistic. **C.** Calculated rate of evolution for viruses in clade B. For the purposes of calculating a rate of evolution the first sample collected from the patient was included as part of clade B. **D.** Estimation of N_e_ for clade B. The results of simulations shown here are identical to those in part B of the figure.

A simulation based analysis, measuring the extent of evolution in idealised Wright-Fisher populations^13^, inferred an effective population size of 2.5 × 10^7^ (95% confidence range 1.0 × 10^7^ to 9.0 × 10^7^) for viruses in clade A before the use of favipiravir(Fig. 2B). This value is substantially larger than estimates made recently for within-host HIV infection^33,34^, and suggests that even weak selection could easily overcome genetic drift. Data from clade B gave a lower estimated value of 2 × 10^6^, (95% confidence range 4 × 10^5^ to 2 × 10^8^) perhaps reflecting the less frequent observation of samples in that clade (Fig. 2C, D).

The value of Ne might be lowered by population structure within the influenza infection^35^. The partial onset of zanamivir resistant alleles^36^, sporadically observed at intermediate frequency after the administration of the drug (Fig. 2S1), is suggestive of population structure going beyond our simple division into clades; the random sampling of viruses from resistant and susceptible subpopulations would produce this behaviour.

Positive selection might have led us to underestimate N_e_. While viral evolution was generally not driven by positive selection (Fig. 2S2), any such selection (e.g. for zanamivir resistance) would increase the rate of viral evolution, lowering our inferred value. Despite this, our result is clear. Once an infection is established, selection will dominate the stochastic effects of drift upon within-host evolution.

The dataset we considered is particularly suited to our calculation. The long period of infection combined with frequent sampling allowed for the characterisation of a slow rate of evolution amidst population structure and noise in the data. Further, the absence of strong selection reduced the error intrinsic to our inference approach, which assumed an idealised neutral population. To provide further validation we repeated our approach on data describing long-term influenza A/H3N2 infection in four immunocompromised adults^37^. The estimates for N_e_ we obtained, of between 3 × 10^5^ and 1 × 10^6^ (Fig. 2S3), while high, were smaller than for our flu B case, potentially being reduced by an increased influence of selection.

We believe that our study provides a first realistic estimate of within-host effective population size for severe influenza infection in humans. The viral load in the influenza B case was high, representative of hospitalised cases of childhood influenza infection. However, the magnitude of our inferred effective size, of order 10^7^, suggests that selection will predominate even at lower viral loads. Our result supports the idea that the observed lack of within-host variation in typical cases of influenza^6,7^ can be explained by the short period of infection; the stochastic effects of genetic drift do not limit the impact of positive selection. Rather, as influenza infections are founded by small numbers of viral particles^7,38^, the majority of low-frequency variants must arise through *de novo* mutation. In a typical infection, very strong selection is required for such variants to reach a substantial frequency in the population^39^. We suggest that, while not being confounded by drift, selection does not usually have time to fix novel variants in the population. Clinical evidence suggests that in cases of longer infection, or in the emergence of antiviral resistance, selection does influence the evolutionary outcome of infection^37,40–44^.

Our result highlights the potential importance of longer infections in the adaptation of global influenza populations, particularly where some adaptive immune response remains. A newly emergent variant under strong positive selection increases faster than linearly in frequency^45^. Given a large N_e_, implying efficient selection, additional days of infection will have a disproportionate influence upon the potential transmission of adaptive variants. This does not imply that longer infections are the sole driving force behind global viral adaptation^37^; selective effects affecting viral transmissibility^30^ would provide an alternative explanation. However, our work suggests that longer-term infections may be an important area of study in the quest to better understand global influenza virus evolution.

## Methods

### Summary

In a single-locus haploid system, the expected change in a variant allele with frequency q caused by genetic drift is given by the formula^46^

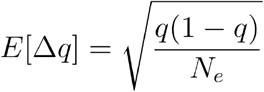

This fact has been exploited to evaluate the size of transmission bottlenecks in influenza infection, comparing statistics of genome sequence data collected before and after a transmission event^30,47^. Such a calculation may be affected by noise in the sequencing of a population, particularly where the extent of noise outweighs the genuine change in a population^30^. Noise in sequence data may be caused by unrepresentative sampling of a population or by error in the experimental process itself^29^.

Because of noise, the comparison of two sequence samples is not a good way of establishing Ne if this statistic may be high. Here we look at an alternative statistic, namely the sum change in variant frequencies in a population; this statistic describes the rate of evolution of the population as a whole. We apply a method of linear regression to multiple samples from the population to establish this rate. We then use simulations to identify the value of N_e_ that, given the diversity of the population, reproduces this rate of evolutionary change. Our approach is robust to noise in the data, the regression calculation identifying the underlying rate of the evolution of the population rather than the simple observed distance between samples.

### Sequence data and bioinformatics

Sequence data describing the evolution of the infection was generated as part of a previous study^32^. Data, edited to remove human genome sequence data, have been deposited in the Sequence Read Archive with BioProject ID PRJNA601176. The HCV data used in validating the sequencing pipeline (see below) was previously deposited in the Sequence Read Archive with BioProject ID PRJNA380188. Processed files describing raw variant frequencies for both datasets are available, along with code used in this project, at https://github.com/cjri/FluBData.

Short-read data were aligned first to a broad set of influenza sequences. Sequences from this set to which the highest number of reads aligned were identified and used to carry out a second short-read alignment. The SAMFIRE software package was then used to filter the short-read data with a PHRED score cutoff of 30, to identify consensus sequences, and to calculate the number of each nucleotide found at each position in the genome. SAMFIRE is available from https://github.com/cjri/samfire.

### Calculation of evolutionary rates

Variant frequencies at different time points during infection were used to calculate a rate of change in the population over time. For each locus in the genome for which data were collected, we calculated the nucleotide frequencies, describing the frequency at each locus *i* of each of the nucleotides at the time of sampling *t*. We then calculated differences in the frequencies observed at each locus *i* using a generalisation of the Hamming distance

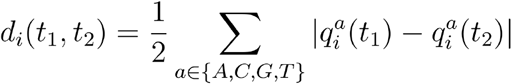

where the term inside the sum indicates the absolute difference between the frequency of allele *a* at locus *i*. The statistic *d_i_* is equal to one in the case of a substitution, for example where only A nucleotides are observed in one sample and only G nucleotides in another. However, in contrast to the Hamming distance it further captures smaller changes in allele frequencies, lesser changes producing values between zero and one, such that a change of a variant frequency from 45% to 55% at a two-allele locus would equate to a distance of 0.1, representing half of the sum of the absolute changes in each of the two frequencies.

Having calculated *d_i_* statistics for each locus, the total distance between two samples was calculated as

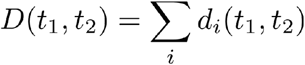

where the sum over *i* was conducted over all loci in the viral genome.

In order to calculate a rate of evolution, between-sample distances were plotted against the separation in time between samples. Linear regression was performed using the Mathematica 11 software package, using the same package to calculate a 97.5% confidence interval for the result. In this case the gradient of the linear model gives the rate of change in time of the genetic distance between samples, averaged across the dataset, providing an estimate of the rate of evolution of the population. The intersection of the line with the vertical axis, equal to the nominal distance between samples at time zero, gives an indication of the extent of ‘noise’ in the data, which may arise from artefacts in either the sampling or sequencing of viruses from the host^29^.

Calculation of synonymous and non-synonymous rates of evolution in the population were calculated in the same manner, with the exception that the distances *d_i_* were calculated over individual nucleotides rather than in a per-locus manner. We calculate

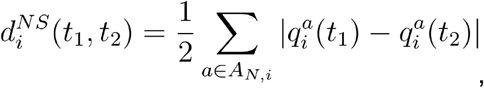

and

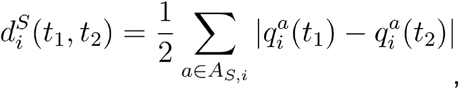

where A_N,i_ and A_S,i_ are the sets of nucleotides at position *i* in the genome which respectively induce non-synonymous and synonymous changes in the consensus sequence. Synonymous and non-synonymous variants were identified with respect to influenza B protein sequences; a nucleotide substitution was defined as being non-synonymous if it induced a change in the coded protein in at least one viral protein sequence. Mean rates of synonymous and non-synonymous evolution were expressed as mean values per nucleotide, reflecting the differing numbers of each type of potential substitution in the viral genome.

### Estimating the effective population size: Wright-Fisher simulation

We used a Wright-Fisher model to simulate the evolution of viral populations, identifying the population size that gave an equivalent rate of evolution to the real data^13^. Data from the viral population gives an estimated rate of evolution per day whereas a Wright-Fisher simulation gives an estimated rate of evolution per generation. We therefore scaled the former to match the experimentally ascertained estimate of 10 hours per generation for influenza B^53^.

To conduct our simulation we constructed a population of N viruses. Each simulated virus had a genome comprised of eight segments, each identical in length to the corresponding segment of the influenza B virus sampled from the patient. The genetic composition of the viral population at the beginning of the simulation was determined by the observed frequencies of non-consensus alleles in a random sample collected from the population. At each locus, a multinomial sample of viruses were chosen to be assigned each of the non-consensus alleles in accordance with the observed frequencies. Variant alleles were assigned independently for each locus, with no intrinsic association between alleles. The sample collected on 23rd November 2017 was excluded as a starting point from this analysis due to its low read depth; all other samples had a mean read depth in excess of 2000-fold coverage.

We simulated a single generation of the evolution of our population under genetic drift, generating a random sample of N viruses from the population. We calculated allele frequency data from the initial and final populations, using these to calculate the distance in sequence space through which the population had evolved according to the modified Hamming distance described above.

For each population size tested, our simulation was run 400 times, using the data to produce a 97.5% confidence interval for the extent of evolutionary change at a given effective population size. The extent of evolution of the real population was then compared to the results from our simulated populations, giving an inference of the effective size of the viral population.

Amendments were made to this basic approach

### Accounting for false-positive variants in sequencing: Estimating a false positive rate

The evolutionary distance calculated by our method is dependent upon the extent of diversity in the viral population. Given a greater number of polymorphic alleles in a system, the evolutionary distance, calculated as the sum of allele frequency changes, will also increase. While the experimental pipeline we used has been shown to perform well in capturing within-host viral diversity^54^, the possibility remains that sequencing could contribute additional diversity to the initial populations used in our simulation. We therefore made an estimate of the extent to which our sequencing process led to the false identification of variants.

To achieve this we used data from a previous study describing the repeat sequencing of hepatitis C virus (HCV) samples from a host^29^; data in this previous study were collected using the same sequencing pipeline as that used to collect the data considered here and therefore provide a generic measure of the level of false positive variation. The data we analysed, coded as HCV01 in the original study, comprised four clinical HCV samples, each of which was split following nucleic acid extraction. Some replicate samples were processed using a DNase depletion method before all samples went through cDNA synthesis, library preparation and sequencing. DNase depletion led to samples with lower read depth; we here compared sequence data collected from the non-depleted replicates of each sample. Variant frequencies within this dataset, where variation was observed in more than one sample, are shown in Fig. 2S4.

Considering the real viral sample, we note that at any given genetic locus, a minority variant either exists or does not exist according to some well-defined criterion. (For the moment the way in which variation is defined is not important; methods for defining variation, which include the use of a frequency threshold, are discussed later.) We denote the possible states of a locus as P and N, according to whether the locus is positive or negative for variation. We suppose that the probability that a random locus in the genome has a minority variant is given by P_P_, leading to the equivalent statistic P_N_ = 1-P_P_.

Sequencing of a specific position in the genome results in the observation or non-observation of a variant. In our data we have sets of two replicate observations of each position in the genome, giving for each minority variant the possible outcomes VV, VX, XV, and XX, where V corresponds to the observation of a variant, and × corresponds to the non-observation of a variant. These observations contain errors; we denote the true positive, false positive, true negative and false negative rates of the variant identification process by P_V|P_, P_V|N_, P_X|N_, and P_X|P_ respectively. In this notation, V|P indicates the observation of a variant conditional on the variant being a true positive.

The underlying purpose of our calculation is to remove falsely detected variation from the population. We begin by assuming that the false negative rate of detecting variants is equal to zero. That is, where we do not see a variant in the sequence data, we assume that a variant is never actually present. This is a conservative step in so far as we never add unobserved variation to the population. Our assumption gives the result that the false negative rate, P_X|P_ = 0. In so far that a variant is never unobserved it follows that the true positive rate P_V|P_ = 1.

We may now construct expressions for the probabilities of observing each of the four possible outcomes. Noting that P_V|N_ + P_X|N_ = 1. we obtain

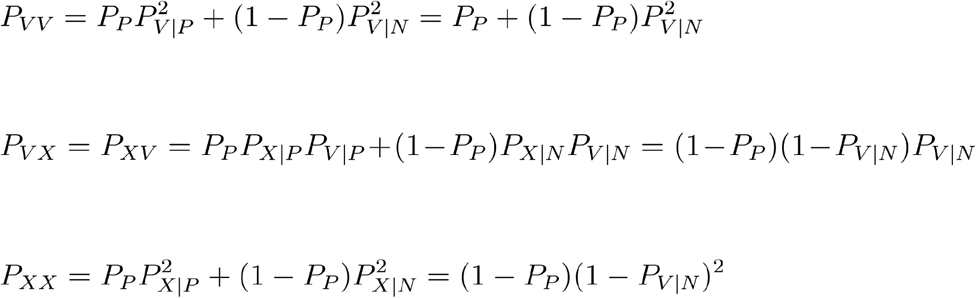

Thus the outcome probabilities may be expressed in terms of the underlying probability of a position having a variant, P_P_, and the false positive rate P_V|N_.

We next processed our sequence replicate data, considering only sites that were sequenced to a read depth of at least 2000-fold coverage. For each locus in a dataset, we calculated the observed frequency of each of the nucleotides A, C, G, and T, generating pairs which described these frequencies in each of our two replicate datasets. Removing pairs in which an allele has a frequency of more than 0.5 in either of the two datasets, we obtained a list of minority variants from each locus, generally comprising three allele frequency pairs per locus. If it is correct that two of the three minority alleles have very low frequencies, the frequencies are close to being statistically independent; the existence of a very few alleles of one minority type does not greatly affect the probability of another variant allele being observed in another read. We note that, of the more than 73 thousand sites sequenced, only 56, fewer than 0.1%, had more than one minority variant at a frequency greater than 1%. We proceeded on the assumption that each pair of minority frequencies was statistically independent of the others.

From the repeated observations of sites, we may count the number of observations of each of the four outcomes; given a total of N pairs we denote these as N_VV_, N_VX_, N_XV_, and N_XX_. Under our model of independent pairs we constructed the multinomial log likelihood of the underlying variant and false positive rates.

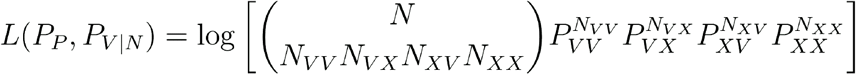

where the terms P_ab_ are constructed from P_P_ and P_V|N_ according to the equations above.

Given a set of paired observations, we calculated the maximum likelihood values of P_P_ and P_V|N_. From these statistics we are able to calculate the positive predictive value of sequencing, namely the proportion of observed variants that are true positives. This is achieved by dividing the probability that a true positive was detected (as P_V|P_ = 1, equal to the probability that a locus has a minority variant), by the probability that a variant was detected:

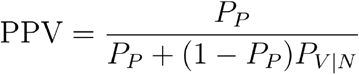

### Frequency dependence of false-positive variant calling

Within our data, our expectation was that minority variants at higher allele frequencies would be more likely to be observed as variants in both replicate samples. We note that, where a frequency cutoff is applied to identify variants, care is required in the above protocol. For example, if a hard threshold was applied, in which variants were called at 1% frequency, a variant that was detected at frequencies of 1.01% and 0.99% would be regarded as having been observed in one case, and not observed in the other, although it likely represents a consistent observation.

In order to assess the frequency dependence of our true positive rate we defined minimum and maximum variant frequency thresholds q^min^ and q^max^, and denoted the replicate observations of a minority variant frequency as q^A^ and q^B^ in the two samples. We further defined the frequency q^cut^ according to the formula

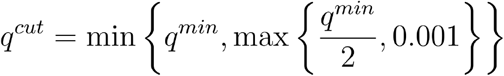

We then defined regions of frequency space as follows:

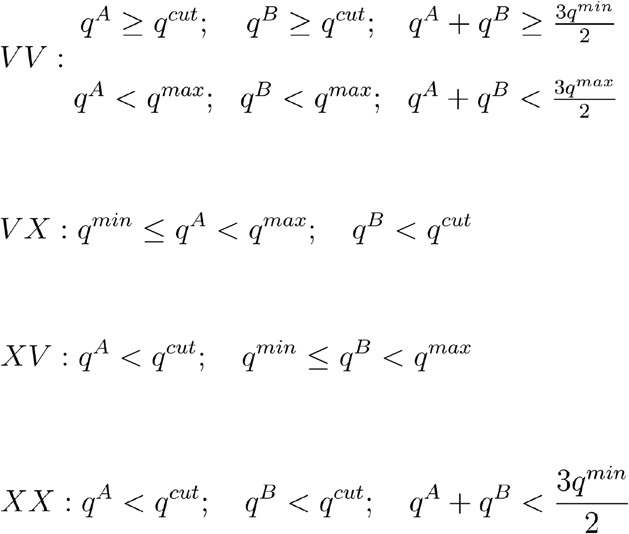

These inequalities are illustrated in Fig. 2S5.

In the above, q^cut^ functions to slightly harshen the criteria for detecting variants at low frequencies. If a variant is observed in one sample at frequency greater than q^min^, then if q^min^ is greater than 0.2%, the frequency in the second sample had to be at least half q^min^ to be counted. If q^min^ was between 0.1% and 0.2%, the frequency in the second sample had to be at least 0.1%, while if q^min^ was less than 0.1%, the frequency in the second sample had to be at least q^min^.

For different ranges of frequency values, q^min^ and q^max^, the proportion of observed variants that were true positives was calculated according to the maximum likelihood method above, using these categorisations. Results are shown in Fig. 2S6. In the process of setting up the initial state of our Wright-Fisher simulation variants observed in the sequence data were considered in turn, drawing a Bernoulli random variable for each variant. Variants were included in the initial simulated population with probability equal to the proportion of observed variants that were estimated to be true positives.

### Accounting for mutation-selection balance

To account for our neglect of mutation, a frequency cutoff was applied to our simulation data. Under a pure process of genetic drift, low-frequency variants in our population are likely to die out, reaching a frequency of zero. In a real population, this would not occur, variants being sustained at low frequencies by a balance of mutation and purifying selection^55,56^. To correct for this we post-processed the initial and final frequency values from our simulations before calculating our distance, imposing a minimum minority allele frequency of 0.1%. All changes in allele frequency below this threshold were ignored, such that, for example, if a variant changed from 0.5% to 0%, this was processed after the fact so that the variant changed from 0.5% to 0.1%. The choice of threshold here is conservative; leading to a conservatively low estimate of N_e_.

### Confidence intervals

Confidence intervals for the effective population size were calculated as the overlap of 97.5% confidence intervals for the evolutionary rates in the observed data, calculated from the regression for the real data, and estimated from the simulated statistics. The overlap of these values gives an approximate 95% confidence interval for N_e_.

### Approximations in the Wright-Fisher model

In the calculation to set up an initial viral population, the assignment of minority alleles to sequences becomes slow at large population sizes. Our code simulated viral genomes; a variant allele was included into the population by choosing an appropriate proportion of genomes to which the variant was assigned. For greater computational efficiency we used a pseudo-random approach for choosing genomes. Given a population size N, we generated a set P of prime numbers that were each larger than N. Given some desired allele frequency q we wish to choose qN genomes to which to assign the variant. We therefore calculated the set of numbers

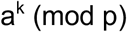

where p is a prime number sampled at random from the set P, and a is a randomly chosen primitive root of p. Given this choice of a and p, the values a^k^ (where k is an integer between 1 and p-1) form a pseudorandom permutation of the numbers from 1 to p-1. We constructed a set of qN genomes by choosing genomes indexed in turn by the elements of this set, beginning from k=1, and discarding values greater than N.

To achieve calculations for population sizes larger than 10^7^ we implemented a statistical averaging method. We generated a single population of size 10^6^, then generated 200 outcomes of a single generation of the same size, recording allele frequencies in each case. In order to simulate a value of N of size r × 10^6^ we compared the frequencies of the initial population to the mean frequencies of a random set of r outcomes. This is equivalent of simulating transmission from a population of size r × 10^6^ in which the initial population contains r copies of each of one of 10^6^ genotypes.

### Phylogenetic analysis

Consensus sequences of data were analysed using the BEAST2 software package^48^. Consensus sequences from each viral segment were concatenated then aligned using MUSCLE^49^ before performing a phylogenetic analysis on the whole genome sequence alignment. The B/Venezuela/02/2016 sequence was used to root the alignment, the haemagglutinin segment of this virus having been identified as being very close to those from the patient. Trees were generated using the HKY substitution model^50^. A Monte Carlo process was run for 10 million iterations, generating a consensus tree with TreeAnnotator using the first 10% of trees as burn-in. Figures were made using the FigTree package (http://tree.bio.ed.ac.uk/software/figtree/).

### Haplotype reconstruction

Haplotype reconstruction was performed using multi-locus polymorphism data generated by the SAMFIRE software package^51^. Variant loci in the genome were identified as those at which a change in the consensus nucleotide was observed between the initial and the final consensus. The short-read data were then processed, converting reads into strings of alleles observed at these loci; a single paired-end read may describe alleles at none, one, or multiple loci. Next, these strings were combined using a combinatorial algorithm to construct a list of single-segment haplotypes, sufficient to explain all of the observed data; no frequencies were inferred at this point. Finally, a Dirichlet-multinomial model was used to infer the maximum likelihood frequencies of each haplotype given the data from each time point^52^. Formally, we divided reads into sets, according to the loci at which they described alleles. A multi-locus variant consists of an observation of some specific alleles at the loci in question. By way of notation, we denote by the number of reads in set *i* which describe the multi-locus variant *a*, and denote the total number of reads in the set as *N_i_*. Given a set of haplotypes with frequencies given by the elements of the vector ****q****, we write as the summed frequencies of haplotypes that match each multi-locus variant *a* in set *i*. For example, the haplotypes ATA and ATG would both match the multi-locus variant AT-describing alleles at only the first two loci. We now express a likelihood for the haplotype frequencies:

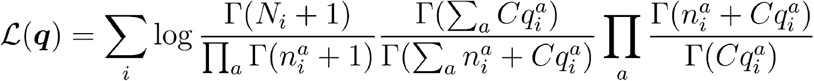

Here the parameter C describes the extent of noise in the sequence data, a lower value indicating a lower confidence in the sequence data. Haplotype reconstruction was performed by finding the maximum likelihood value of the vector of haplotype frequencies **q**. A value of C=200 was chosen for the calculation, representing a conservative estimate given the prior performance of the sequencing pipeline used in this study^29^. In contrast to previous calculations in which an evolutionary model was fitted to data^52^, haplotype frequencies for each time point and for each viral segment were in this case inferred independently, with no underlying evolutionary model.

### Data describing influenza A/H3N2 infection

Our analysis of data describing long-term influenza A/H3N2 infection was performed on data from a previous study^37^. As our method does not require an exceptional quality of sequencing data to calculate a rate of evolution more samples were included in our analysis than were examined in the original study. Using the codes established in the previous study, we used samples from patient W from days 0, 7, 14, 21, 28, 56, 62, 67 and 76; from patient X from days 0, 7, 14, 21, 28, 42, and 72; from patient Y from days 0, 7, 14, 21, 28, 35, 48, 56, and 70; from patient Z from days 14, 15, 20, 25, 41, 48, 55, 62, and 69. An identical procedure to that used to estimate Ne from the influenza B data was applied, calculating a rate of evolution per day from sequence data, scaling this to a rate per generation (in this case a seven hour generation time was modelled^53^), and then running simulations to estimate N_e_. We note that the estimates of false positive rate generated for the influenza B data were applied equally in this case, due to not having equivalent data to re-estimate these values. Examining the data from patient W, our distance measurements suggested potential population structure involving the samples collected on days 62 and 69; these samples were excluded from our regression analysis.

---

**Figure 1 supplement 1.**
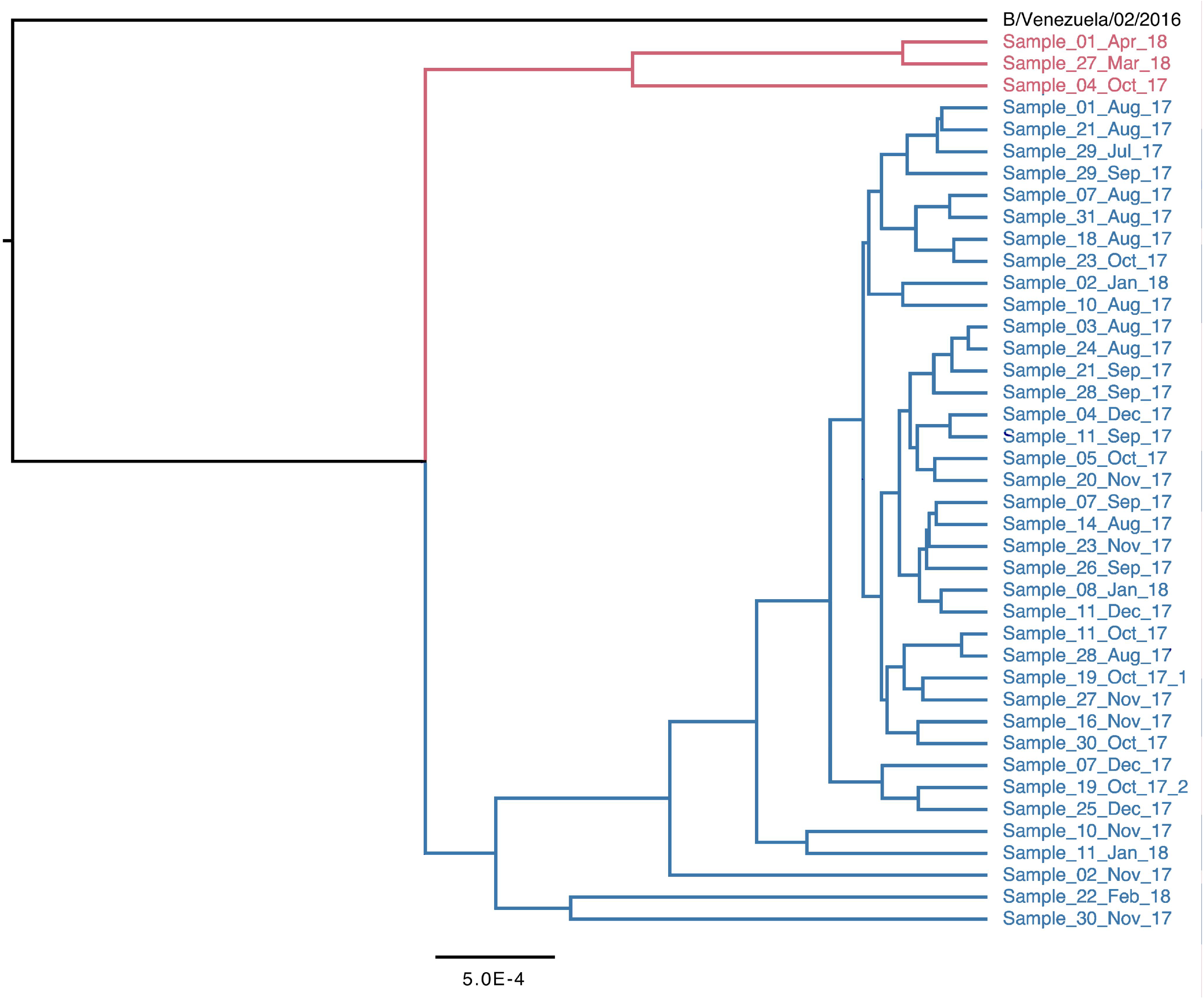
Complete phylogeny of whole-genome viral consensus sequences, coloured by clade.

**Figure 1 supplement 2:**
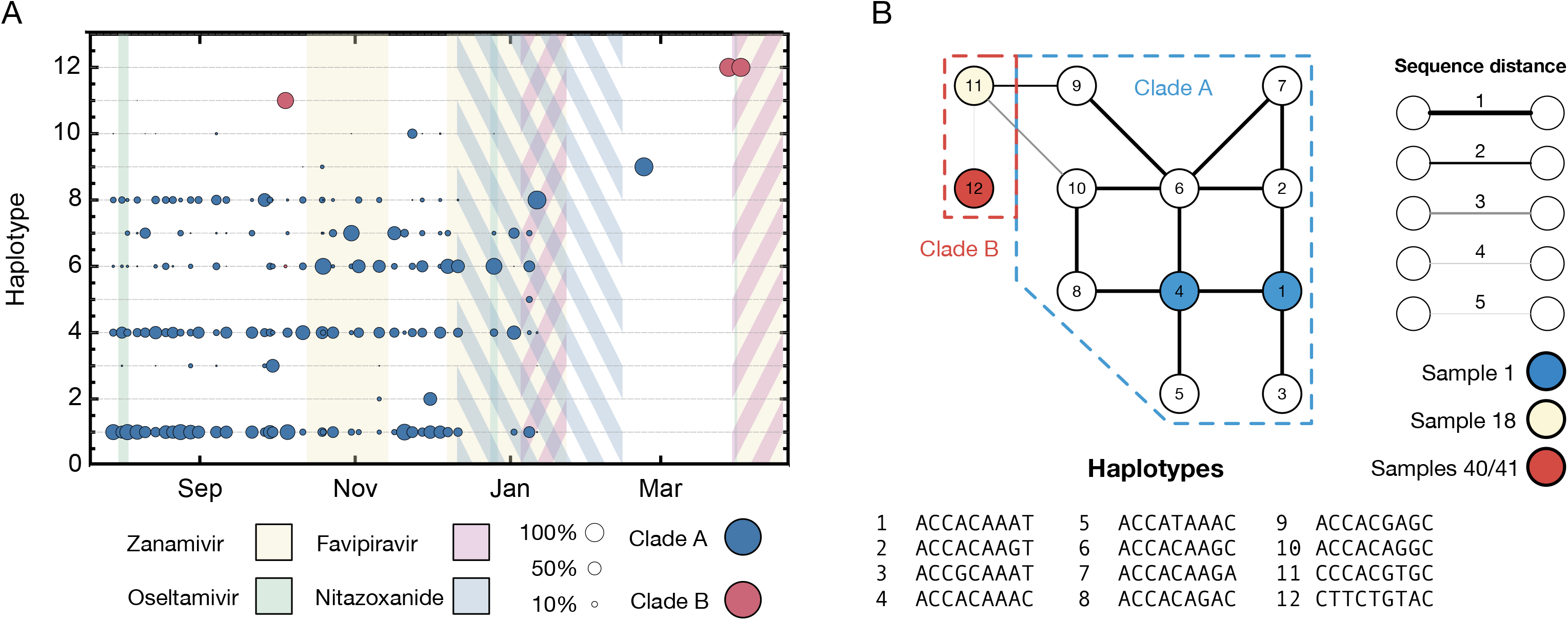
**A.** Sub-consensus structure of the viral population inferred via a haplotype reconstruction algorithm using data from the haemagglutinin segment. A division of sequences into two clades is visible, with samples including largely distinct viral genotypes. The area of each circle is proportional to the amount of virus in each clade. Haplotypes reaching a frequency of at least 10% in at least one time point are shown. Haplotypes spanned the loci 258, 261, 364, 451, 521, 541, 635, and 641 in the HA segment. **B.** Evolutionary relationship between the haplotypes; clade B is distinct from and evolves away from those sequences comprising the initial infection. Numbers refer to the distinct haplotypes identified within the population.

**Figure 2 supplement 1:**
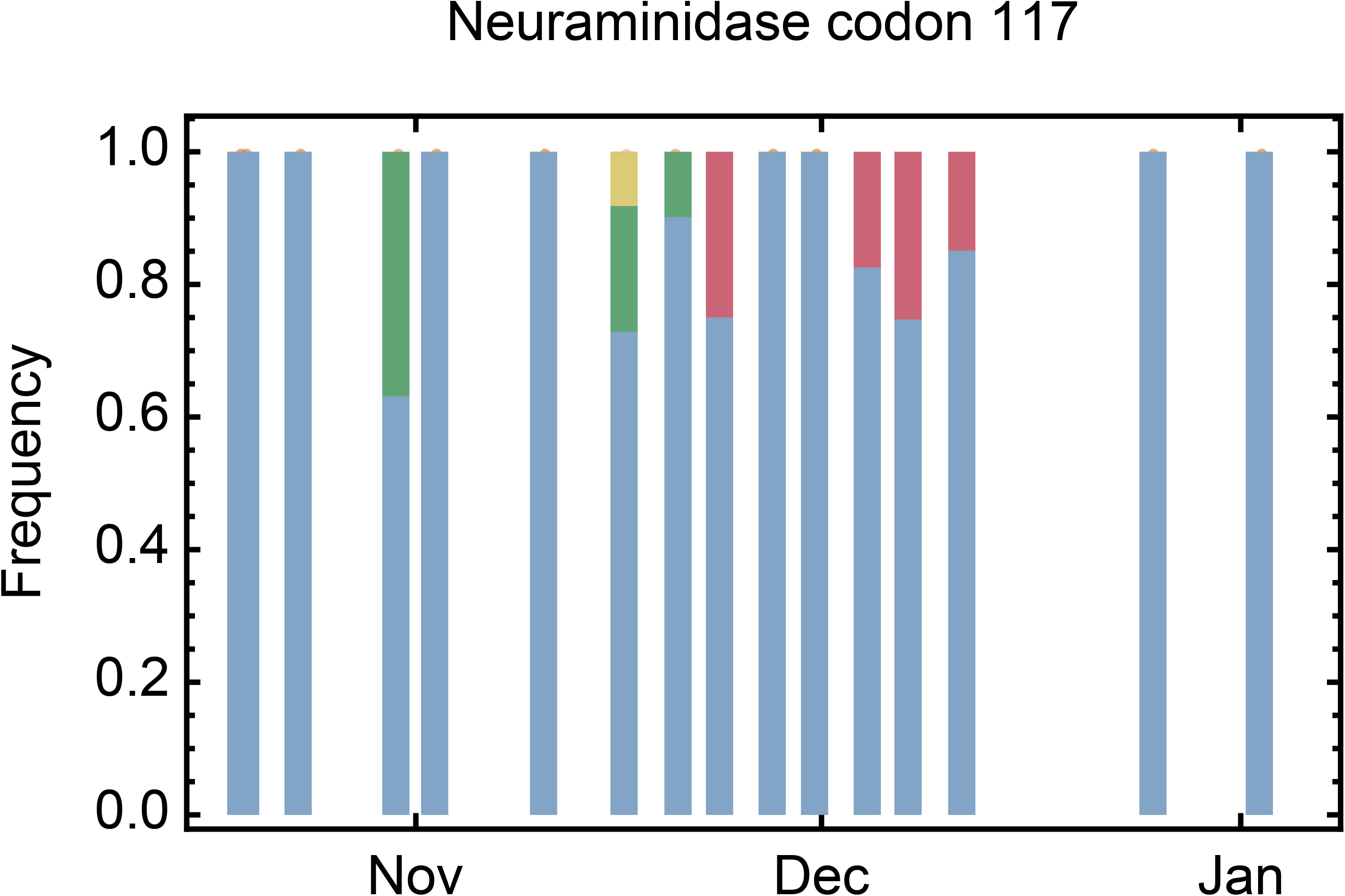
Amino acids present at codon 117 of the neuramindase segment of the virus after the first administration of zanamivir. The consensus glutamate nucleotide (blue) was sometimes replaced by glycine (green), valine (yellow), and alanine (red). Glycine and alanine are associated with zanamivir resistance in influenza B.

**Figure 2 supplement 2:**
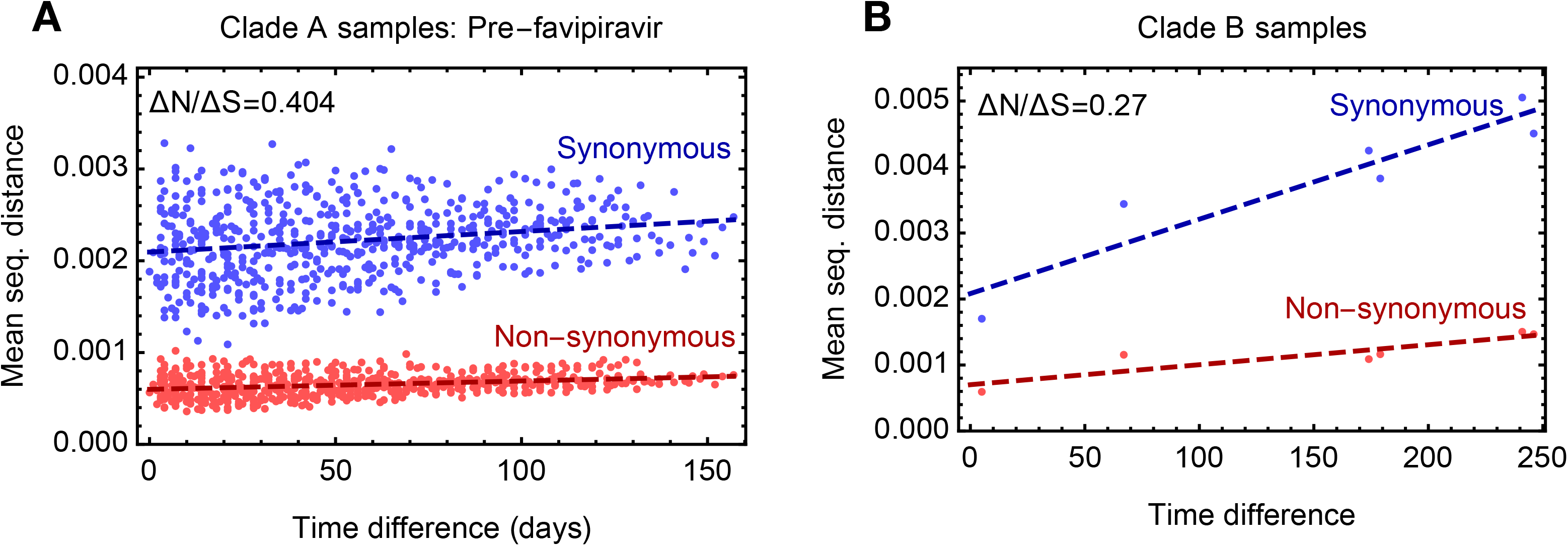
**A.** Comparison of rates of synonymous and non-synonymous evolution for viruses in clade A up to the time of the administration of favipiravir. The distance between two samples is calculated as the mean absolute difference in allele frequency, as averaged over synonymous and non-synonymous positions in the genome. B. Comparison of rates of synonymous and non-synonymous evolution for viruses in clade B. The rate of evolution in both clades was slower at non-synonymous sites than at synonymous sites, suggesting a general pattern of purifying selection at non-synonymous sites. Change in the population was not as a whole driven by positive selection.

**Figure 2 supplement 3:**
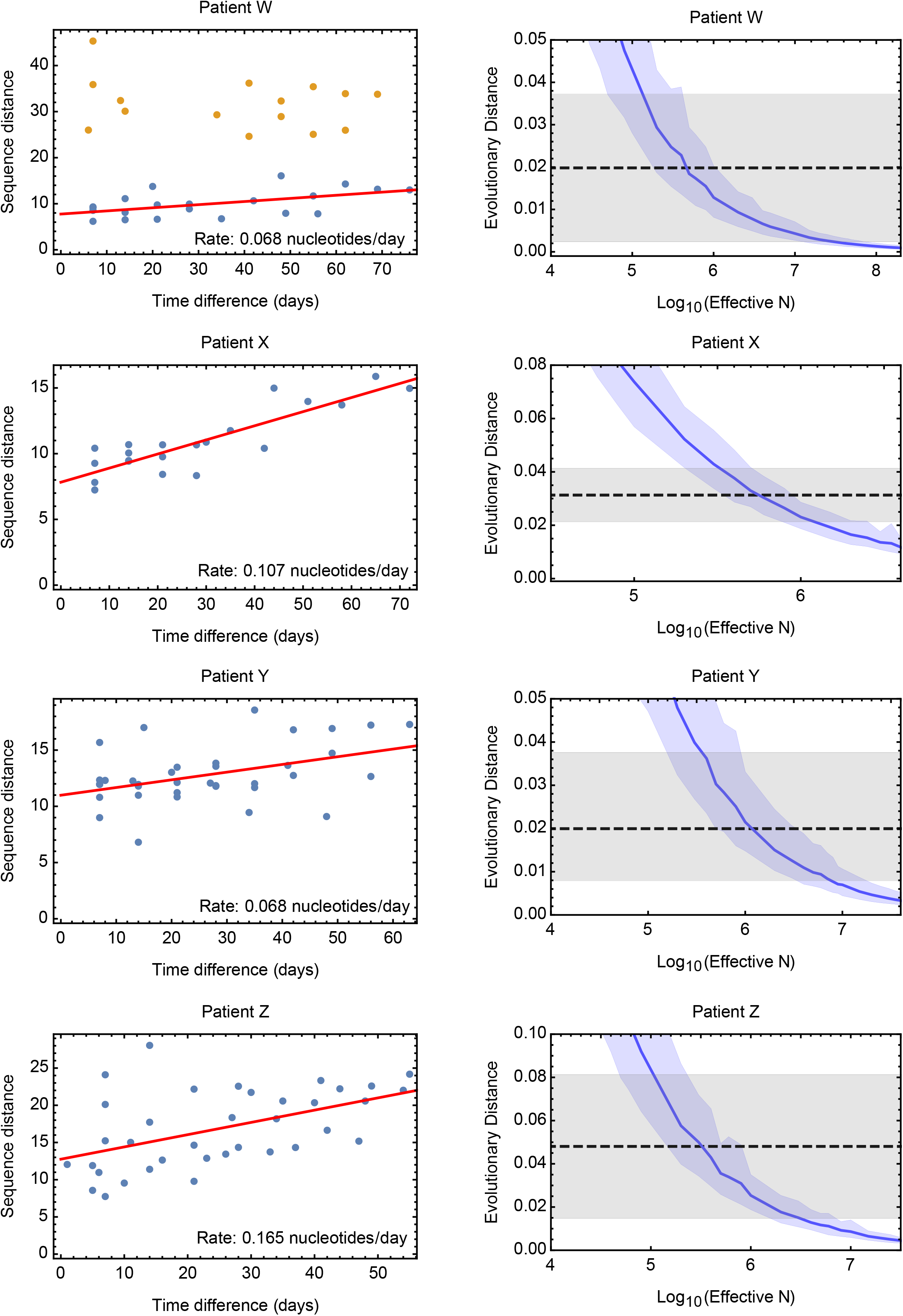
Estimates of the effective population size for data from a study of long-term influenza A/H3N2 infection in four patients. Patients are denoted with the letters assigned them in the original study^37^. Rates of evolution within each patient were calculated by linear regression, conducted on a plot of evolutionary versus temporal distance between samples. The inferred regression line is shown in red for each dataset. For Patient W samples collected at two time points appear as outliers in the distance plot; distances involving these samples, shown in yellow, were excluded from the calculation. Accompanying plots show distances inferred via simulation compared to the inferred rates. A solid blue line shows the mean, with shading indicating an approximate 97.5% confidence interval around the mean. Statistics were calculated from sets of 400 simulations conducted at each value of N_e_.

**Figure 2 supplement 4:**
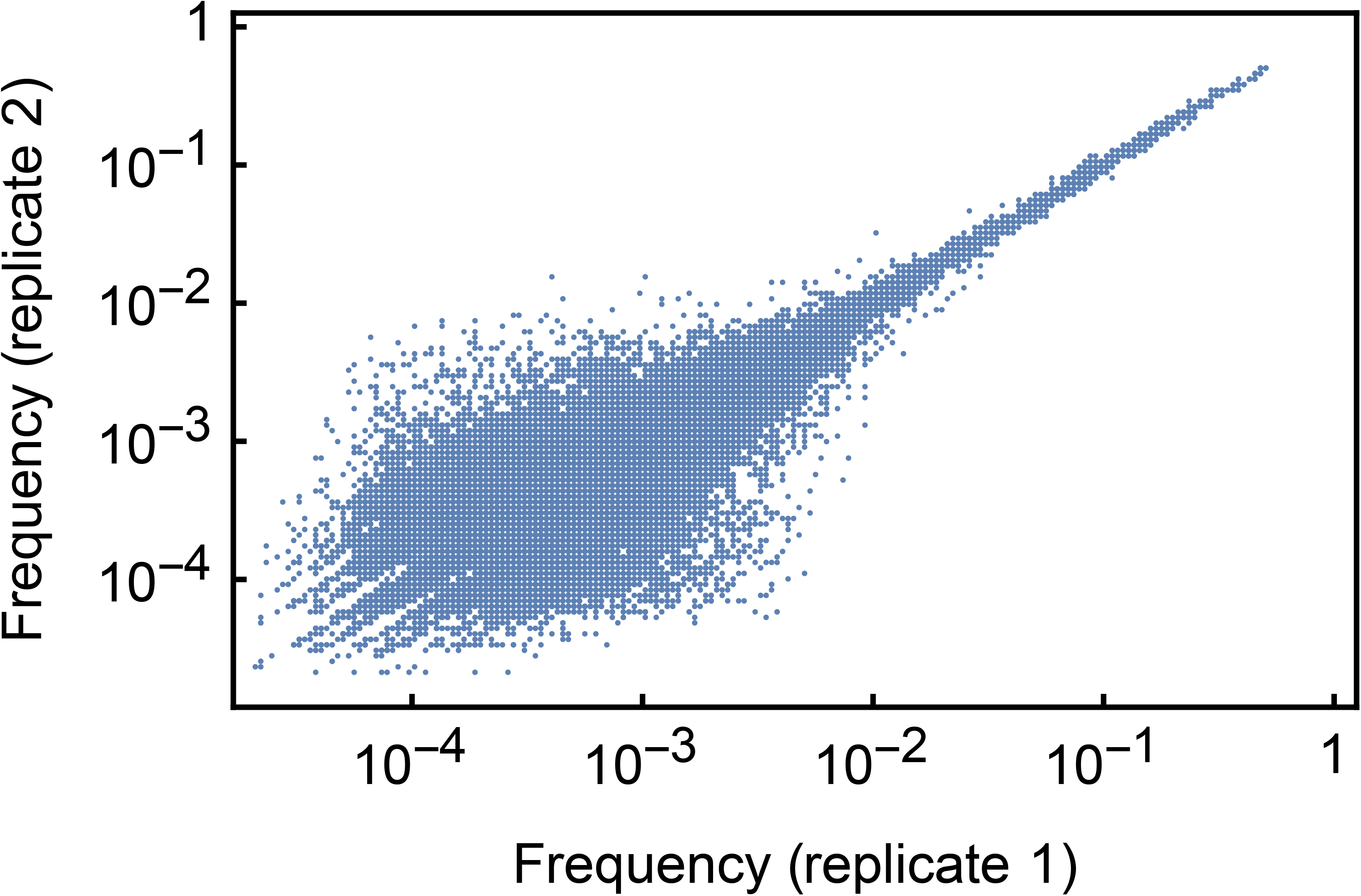
Frequencies of minority variant alleles identified in the HCV01 dataset used to evaluate the accuracy of variant calling in our sequencing pipeline. Samples in this dataset were split following RNA extraction with replicate sets of RNA being processed and sequenced independently. Variants at higher frequencies were identified at more consistent frequencies than variants at lower frequencies.

**Figure 2 supplement 5:**
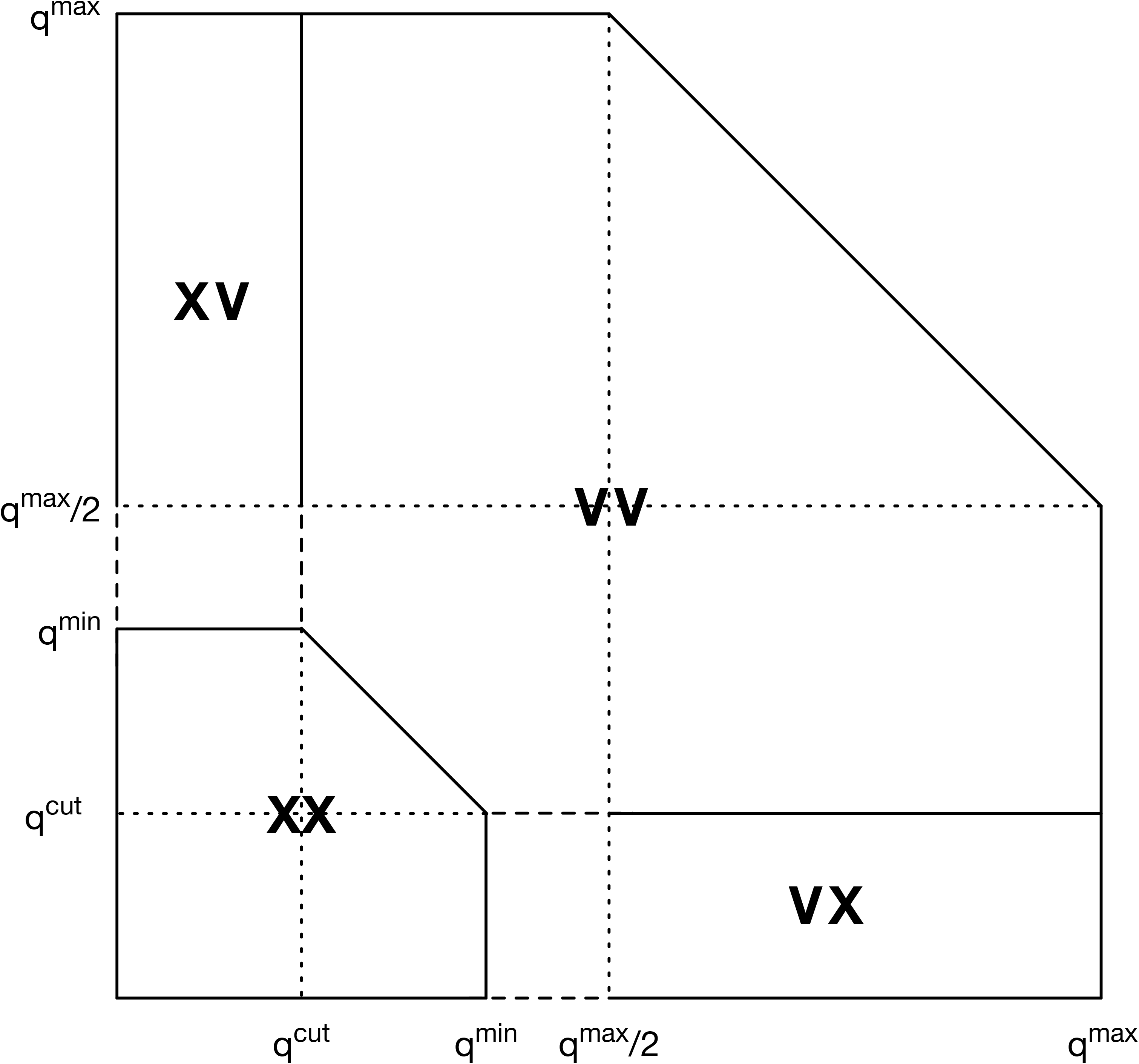
Regions of frequency space used to define observations and non-observations of allele frequencies. V indicates the identification of a variant, while X indicates the non-identification of a variant. Combinations of V and X indicate observations made in two replicate samples.

**Figure 2 supplement 6:**
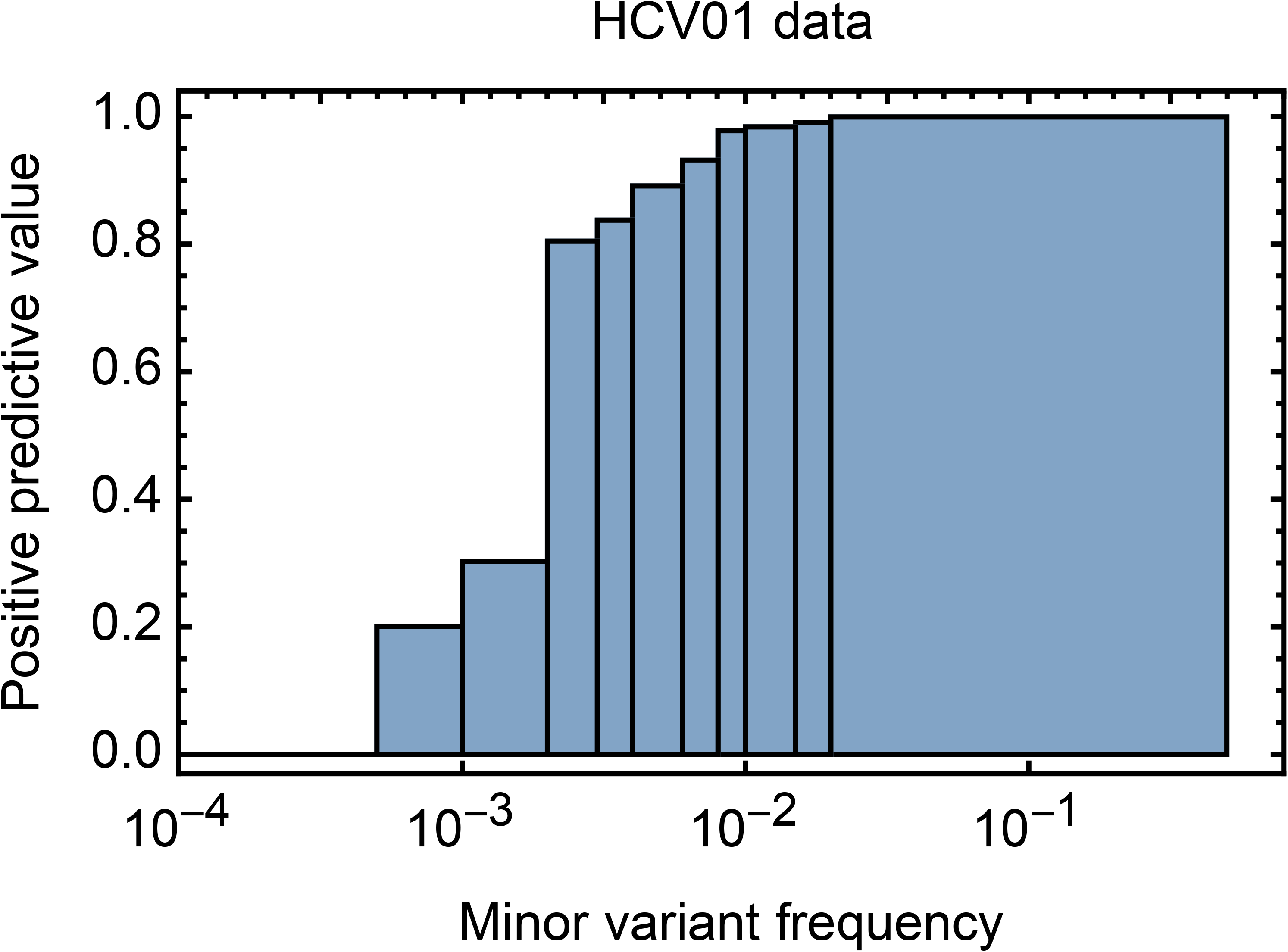
Positive predictive value for minority variants under our sequencing pipeline, calculated at different frequency ranges. While high frequency variants were very reliably identified, the reliability of identifying variants was significantly impaired at lower frequencies.

